# Projecting neurons from lateral entorhinal cortex to basolateral amygdala mediate the encoding of incidental odor-taste associations

**DOI:** 10.1101/2025.01.07.631103

**Authors:** Jose Antonio González-Parra, Vittoria Acciai, Laura Vidal-Palencia, Marc Canela, Arnau Busquets-Garcia

## Abstract

Daily choices are determined by prior direct or indirect associations between low-salience cues and reinforcers. In this study, we used a mouse odor-taste sensory preconditioning task combined with genetic, intersectional and chemogenetic approaches to identify a novel brain circuit involved in mediated learning. We found that neuronal projections from the lateral entorhinal cortex to the basolateral amygdala are engaged during low-salience stimuli associations, which is essential for mediated learning formation.

---

Environmental cues are fundamental in guiding both animal and human daily choices as they evoke innate and conditioned behavioral responses. While classical Pavlovian conditioning has been extensively used to explain most of these behavioral choices, less explored and more complex conditioning processes also dictate such decisions. In this sense, higher-order conditioning has been proposed to account for unexpected aversion or attraction toward stimuli that have never been directly reinforced before^1-3^. Understanding the brain circuits involved in higher-order conditioning is critical for elucidating how the brain encode and store indirect associations between different low-salience stimuli that influence animal behavior. In laboratory settings, rodent sensory preconditioning protocols have proven valuable for studying these intricate cognitive processes^1-6^. In the present study, we combined an odor-taste sensory preconditioning task^1^ with the use of “Targeted Recombination in Active Populations” (TRAP2) mice^7,8^ and intersectional viral approaches to identify and manipulate specific brain circuits through chemogenetic approaches.

Throughout this study, we used an odor-taste sensory preconditioning task adapted from previous findings^1,8^. This task consists in an habituation period to the water deprivation (3 days), a preconditioning phase presenting different odor-taste pairings (Saccharin-Banana and NaCl-Almond) for 6 days, a conditioning phase where one odor (Almond) was devaluated through the injection of Lithium Chloride (LiCl) and, finally, two-choice tests are performed to assess mediated (two-choice test between tastes) or direct (two-choice test between odors) learning (Fig. 1A). Both male and female C57BL6J mice presented mediated and direct learning (Fig. 1B-C) in this adapted paradigm as shown previously^1,8^, with no innate preferences observed between the tastes and odors used (Fig. S1A-C). To study the brain regions involved during the odor-taste associations (i.e. preconditioning phase) and the mediated learning test, we used the TRAP2:Ai14 mouse line that has been extensively applied to identify the brain circuits engaged by diverse behaviors^7,9^. We injected 4-hydroxytamoxifen (4-OHT) just after each preconditioning session (*i*.*e*. to label the activated cells during this phase with tdTomato, in red). 90 minutes after the two-choice test between tastes (mediated learning test), animals were transcardially perfused and c-Fos staining (in green) was performed to identify the brain cells activated during the test and its comparison with cells engaged during preconditioning labelled with tdTomato (Fig. 1D). Male and female TRAP2:Ai14 mouse line showed mediated learning responses as observed with C57BL6J mice (Fig.1E) with no differences in consumption between control (only received water during the protocol) or TRAP (normal odor-taste protocol) group during preconditioning or mediated learning test (Fig. S1D-E). Therefore, we characterized the expression levels of tdTomato and c-Fos in different brain regions classically involved in sensory preconditioning such as the perirhinal^10,11^, the orbitofrontal cortices^12,13^, the hippocampal region^1,14^ and the amygdala^10,15^. Strikingly, whereas no differences were found in other brain regions (Fig. S1F-H), we observed a specific increase of tdTomato positive cells in the basolateral amygdala (BLA) of mice undergoing the preconditioning phase (i.e. odor-taste associations) compared to animals receiving the same amount of water per day (Fig. 1F). In contrast, the number of c-Fos-positive cells remained unchanged between these two experimental groups as observed during the mediated learning task in any of the brain regions analyzed (Fig. 1G and S1F-H). Regarding the colocalization between activated cells during the preconditioning phase and the probe test, we only found a significant decrease in the perirhinal cortex (Fig 1H and S1G). Notably, the presentation of one stimulus alone through the protocol did not increase the neuronal activation in BLA suggesting that BLA activity is only modulated by incidental associations between neutral stimuli (Fig S1I). Overall, these findings pointed to the BLA as a key brain region to encode low-salience stimuli associations in our odor-taste sensory preconditioning paradigm.

**Fig. 1.**
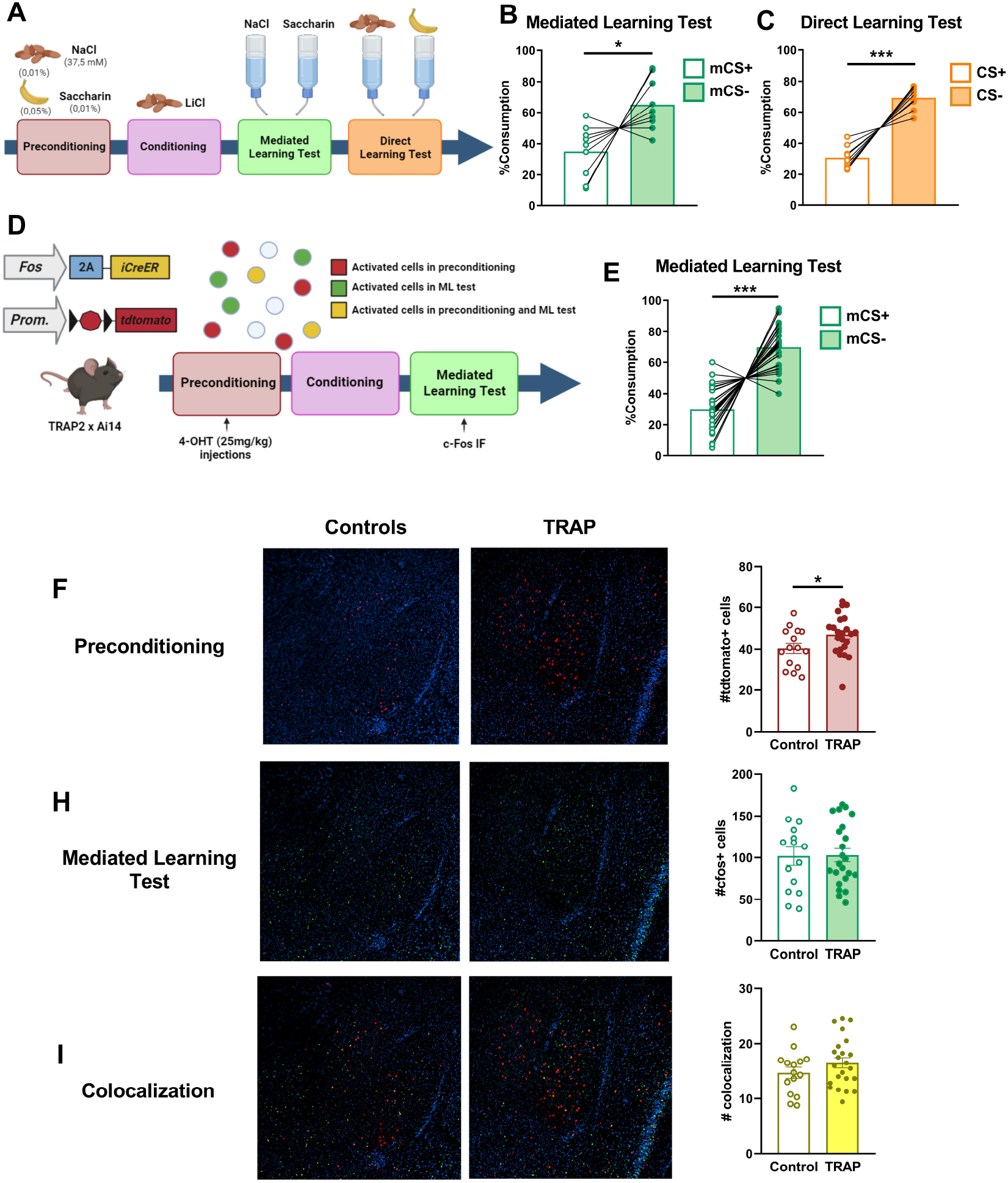
Enhanced Activity in the Basolateral Amygdala (BLA) During Odor-Taste Incidental Associations. (A) Schematic representation of the odor-taste sensory preconditioning paradigm. Percentage of liquid consumption during mediated (B) and direct (C) learning tests in C57BL/6J mice. (D) Schematic of the protocol used in TRAP2:Ai14 mice to investigate brain regions activated during incidental associations (preconditioning phase) and/or retrieval of mediated learning. (E) Percentage of liquid consumption during mediated learning test in TRAP2:Ai14 mice. Quantification of tdTomato-positive cells during the preconditioning phase (F), c-Fos-positive cells during mediated learning test (G) and colocalized tdTomato/c-Fos-positive cells (H) in the BLA of TRAP2:Ai14 mice. Data are represented as mean values and before-after for liquid consumption or as mean ± SEM for image-based quantifications. For statistical details and n, see Supplementary Table 1. Scale Bar: 25 μm. *** p<0.001; * p<0.05. mCS (mediated conditioned stimulus); CS (conditioned stimulus).

To investigate if the activation of the BLA during the preconditioning phase is causally linked to the formation of mediated learning, we infused an adeno-associated viral (AAV) vector carrying the inhibitory Designer Receptors Exclusively Activated by Designer Drugs (DREADD-Gi) under the CAMKII promoter to inhibit the principal neurons in this brain region (Fig. 2A-B). Specifically, the DREADD agonist JHU37160 dihydrochloride (J60, 0.1 mg/kg, *i*.*p*.)^16^ was injected in male and female mice 1 hour before each preconditioning session. We observed that the inhibition of BLA CamkII-positive cells during odor-taste associations completely blocked the mediated learning response (Fig. 2C), while leaving direct learning unaffected (Fig. 2D). Moreover, J60 administration during preconditioning left unaltered the consumption during the preconditioning phase (Suppl. Figure 2A) or during the mediated learning test (Suppl. Figure 2B) but it increased total consumption during direct learning test (Suppl. Figure 2C). Once we identified BLA as a key brain region involved in the formation of mediated learning, we wondered if the same neuronal engrams activated during odor-taste associations in this brain region were involved in the expression of mediated learning. For this reason, we infused a Cre-Dependent AAV-DREADD-Gi into the BLA of TRAP2:Ai14 mice (Fig. 2E-F). Four weeks later, these mice underwent the sensory preconditioning protocol where 4-OHT was injected just after each preconditioning session to specifically express the DREADD-Gi into the preconditioning-induced activated cells exclusively in the BLA. One hour before the mediated and direct learning test, J60 was injected to inhibit the neuronal ensemble engaged during odor-taste associations but the expression of mediated or direct learning was not affected (Fig. 2G-H and Fig S2 D-F). Overall, these results suggest that a particular neuronal ensemble is activated during odor-taste pairings for the formation of mediated learning although it is not directly involved during the expression of this behavior.

**Fig. 2.**
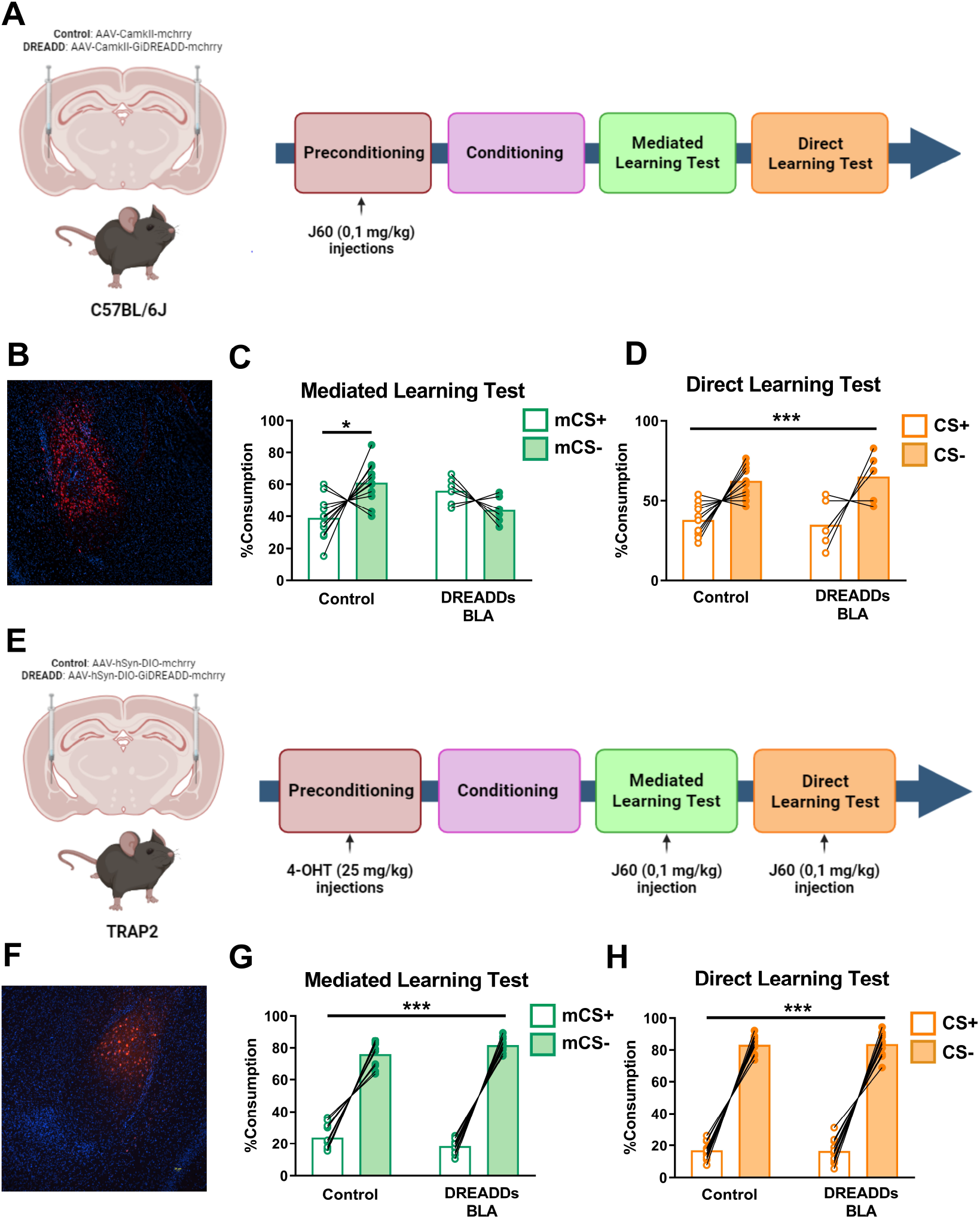
BLA Is Required for Encoding Incidental Associations during odor-taste sensory preconditioning. (A) Schematic illustration of BLA inhibition during the preconditioning phase in C57BL/6J mice. (B) Representative image showing DREADD-Gi expression in the BLA. Percentage of liquid consumption during mediated learning test (C) and direct learning test (D) in C57BL/6J mice. (E) Schematic procedure to inhibit activated preconditioning BLA neurons during mediated learning test in TRAP2:Ai14 mice. (F) Representative image of mCherry expression in the BLA. Percentage of liquid consumption during mediated learning test (G) and direct learning test (H) in TRAP2:Ai14 mice. Data are represented as mean values and before-after for liquid consumption. For statistical details and n, see Supplementary Table 1. Scale Bar: 25 μm. *** p<0.001; * p<0.05. mCS (mediated conditioned stimulus); CS (conditioned stimulus).

The activity of a particular brain region, such as the BLA, could not be uniquely involved in the control of complex behavioral responses such as the encoding of odor-taste pairings. Therefore, it is expected that a complex brain network, involving different brain regions^2^, would be engaged during this preconditioning phase. To identify the BLA inputs being activated during odor-taste pairings, we infused a Cre-Dependent retrograde AAV coupled to the green fluorescent protein GFP into the BLA of male and female TRAP2:Ai14 mice to identify the specific activated neuronal inputs to the BLA (in yellow) during preconditioning (Fig. 3A-C). Four weeks after the stereotaxic surgeries, mice underwent the sensory preconditioning protocol (Fig. 3A-B) and 4-OHT was injected after each preconditioning session. The quantification of cells showing a colocalization between tdTomato (*i*.*e*. cells activated during odor-taste pairings) and GFP (*i*.*e*. neuronal inputs to the BLA) gave us a picture of several brain regions projecting to the BLA that could also participate in the encoding of odor-taste associations. From all these brain regions, the lateral entorhinal (LEnt) cortex appeared to be the brain region with higher neuronal inputs that were activated by odor-taste pairings (Fig. 3C-D and Fig. S3A-D). At this point, we used an intersectional viral approach infusing a retrograde AAV-Cre into the BLA and a Cre-Dependent DREADD-Gi into the LEnt to specifically modulate the activity of neuronal projections from LEnt to BLA (Fig 3E-H). Strikingly, the specific inhibition of these neuronal projections during odor-taste associations by the administration of J60 before each preconditioning session fully blocked the formation of mediated learning (Fig. 3G) without altering the direct learning (Fig. 3H) or the consumption in the different phases (Fig. S3E-G). These results suggest that projecting neurons from LEnt to BLA participate in the encoding of odor-taste associations, being a key brain circuit involved in the formation of mediated learning.

**Fig. 3.**
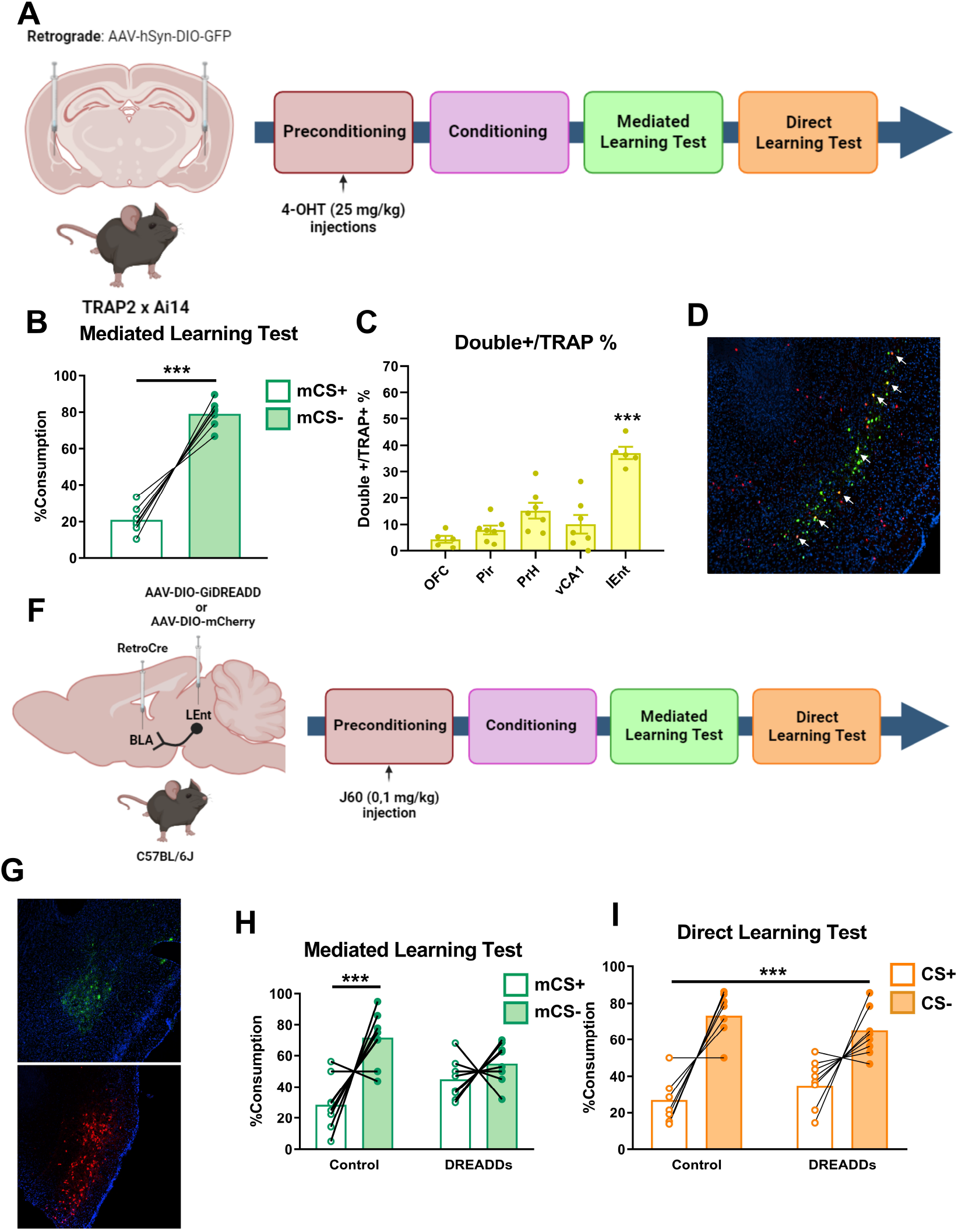
Projections from LEnt to BLA are Critical for the encoding of incidental associations. (A) Schematic procedure to identify activated brain regions projecting to the BLA during preconditioning. (B) Percentage of liquid consumption during mediated learning test. (C) Percentage of activated cells during the preconditioning phase that also project to the BLA (%tdTomato+GFP/tdTomato+) and (D) representative image of the LEnt. (F) Experimental protocol to inactivated LEnt-BLA projection during incidental associations. (G) Representative image showing DREADD-Gi expression in the LEnt and retro-GFP expression in the BLA. Percentage of liquid consumption during mediated learning test (H) and direct learning test (I) in C57BL/6J mice. Data are represented as mean values and before-after for liquid consumption or as mean ± SEM for image-based quantifications. For statistical details and n, see Supplementary Table 1. Scale Bar: 25 μm. White arrows indicate vTdtomato+GFP+ (colocalization) cells*** p<0.001.OFC: Orbitofrontal cortex; Pir: Piriform cortex; PrH: Perirhinal cortex; vCA1: ventral CA1; LEnt: Lateral Entorrinhal cortex. mCS (mediated conditioned stimulus); CS (conditioned stimulus).

The BLA activity has been classically linked to conditioning settings involving aversive reinforcers^17,18^. However, amygdalar circuits have also been involved in novelty and familiarity detection where no aversive salience is present^19-21^, which would be in line with our data showing a particular engagement of this brain region when odor-taste pairings are occurring. In addition, a previous study conducted in rats investigated the involvement of BLA in sensory preconditioning observing that the inhibition of the amygdala was impacting pairings between low-salience stimuli only in a dangerous environment^10^. However, in contrast to our study, the presentation of light and tone was sequential and not simultaneous. Indeed, future studies should clarify the different engagement of brain circuits depending on how the stimuli are presented and how different sensory modalities are encoded. An alternative hypothesis of the role of BLA during the preconditioning phase could be that BLA stores associations between low-salience stimuli to be prepared in case one of these cues are paired with noxious reinforcers in the future and thus anticipating their potential link with aversive outcomes.

Previous findings showed how lesions in the entorhinal cortex impact associative learning such as trace eyeblink conditioning^22^ or trace fear memory^23^. Indeed, the entorhinal cortex has been proposed as a regulatory region in response to different stimuli associations^24,25^. Moreover, the interaction between limbic structures such as the entorhinal cortex and the amygdala are suggested to shape cognitive processes^26,27^. Our study represents one step further in elucidating of the brain circuits involved in mouse sensory preconditioning as it shows the specific involvement of LEnt-BLA projections in the encoding of low-salience stimuli associations and, therefore, in the formation of mediated learning. By deciphering the brain circuits involved in these complex cognitive processes, our work provides new insights that might be helpful to understand brain pathological states such as psychiatric diseases that are accompanied by alterations in mediated learning^8,28^.

## Material and methods

### Animals

Male and female C57BL/6J and *TRAP2:Ai14* mice from 8-14 weeks of age were employed in the study. C57BL/6J were sourced directly from Charles River (Charles Rivers Laboratories, Spain). *TRAP2:Ai14* mice were derived from crosses between homozygous *TRAP2* mice^7^ (Fos^tm2.1(icre/ERT2) Luo^/J; The Jackson Laboratory, Bar Harbor, ME, USA; #030323) and homozygous *Ai14* tdTomato reporter mice (B6.Cg-Gt(ROSA)26Sor^tm14(CAG-tdTomato)Hze^/J; The Jackson Laboratory; #007914). Offspring heterozygous for both transgenes were genotyped following established protocols^7^. Animals were housed in groups of two to four in a specific pathogen-free facility under controlled conditions (temperature: 20-24°C; humidity: 40-70%), with a 12-hour light/dark cycle (lights on at 8 pm, off at 8 am). All mice had continuous access to food and standard environmental enrichment materials, including nesting materials and cardboard houses. Experiments were conducted during the dark phase (9 am to 3 pm), with water access regulated during behavioral testing and provided *ad libitum* before and after testing. Prior to behavioral assessments, mice underwent handling for two consecutive days to acclimate them to the experimenters. All behavioral testing was performed with experimenters blinded to the experimental groups.

This study adhered to the European Directive 2010/63/EU on the protection of animals used for scientific purposes. All procedures received approval from the Animal Ethics Committee of the Parc de Recerca Biomèdica de Barcelona (CEEA-PRBB; approval numbers ABG-19-0055 and ABG-21-0018) and the Generalitat de Catalunya (approval numbers: 10784 and 11590, respectively).

### Drug Preparation and Administration

4-hydroxytamoxifen (4-OHT; Sigma-Aldrich, #H6278) was prepared at a concentration of 20 mg/mL in ethanol by agitating at 37°C for 5 minutes, then aliquoted and stored at –20°C for several weeks. Prior to administration, 4-OHT was dissolved in Kolliphor oil (Merk, Cat # C5135-500G) by shaking at 37°C for 5 min, followed by evaporation of ethanol using a Savant DNA120 SpeedVac concentrator, yielding a final concentration of 10 mg/mL. Saline phosphate buffer (0.1 M) was subsequently added to achieve a final dosing concentration of 25 mg/kg. All 4-OHT injections were administered intraperitoneally (i.p.) immediately after completion of behavioral testing.

J60 dihydrochloride (J60; Hello Bio, # HB6261) was dissolved in sterile saline to reach a concentration of 0,1mg/kg. J60 was intraperitoneally administered 1 hour prior to each preconditioning session and 1 hour prior to both mediated and direct learning tests.

### Chemical Odors and Tastes

For the sensory preconditioning task, solutions were presented in 50-mL drinking tubes placed in the animals’ home cages. Odor solutions included banana (0.05% isoamyl acetate, Sigma-Aldrich #W205532) and almond (0.01% benzaldehyde, Sigma-Aldrich # B1334), while taste solutions consisted of saccharin (0.01%, Sigma-Aldrich, # 109185) and NaCl (37.5 mM, Sigma-Aldrich # 746398). Odor and taste concentrations were selected based on their demonstrated equal preference (Fig. S1B-C)

### Surgical Procedures and Viral Vector Injections

Mice were anesthetized with an i.p injection of ketamine (75mg/kg) and medetomidine (1mg/kg). Analgesia was provided by subcutaneous meloxicam injections (5 mg/kg) administered pre-surgery and continued for two days’ post-surgery. Following anesthesia, the mouse’s head was positioned in a stereotaxic apparatus, and bregma and lambda coordinates were aligned. Small craniotomies were drilled to allow for bilateral infusion of viral vectors at specific coordinates relative to bregma: Basolateral amygdala (BLA): anteroposterior (AP) -1.5 mm, mediolateral (ML) ±3.1 mm, dorsoventral (DV) -5.2 mm; Lateral Entorhinal cortex (LEnt): AP -4.16 mm, ML ±3.7 mm, DV -4.3 mm. To investigate the specific involvement of the BLA during the preconditioning phase, C57BL/6J mice received 300 nL injections of pAAV2-CamkII-hM4D(Gi)-mCherry or pAAV-CaMKIIa-mCherry (Addgene viral prep # 50477-AAV2 and viral prep # 114469-AAV5). For assessing the involvement of BLA activated neurons in preconditioning during the mediated learning test, TRAP2:Ai14 were injected with 250 nL pAAV-hSyn-DIO-hM4D(Gi)-mCherry or pAAV-hSyn-DIO-mCherry (Addgene viral prep #44362-AAV2 and viral prep #50459-AAV2). For studying activity-dependent afferent connections to the BLA during neutral association encoding, TRAP2:Ai14 mice were injected bilaterally with a 250 nL fluorescent retrograde Cre-dependent virus, pAAV-hsyn-DIO-EGFP (Addgene viral prep #50457-AAVrg) bilaterally in the BLA. Additionally, to inhibit projections from the LEnt to the BLA during the preconditioning phase, C57BL/6J mice were co-infused with 250 nL in a proportion 4:1 of retrograde pAAV-EF1a-Cre and retro pAAV-CAG-GFP viral vectors in the BLA (Addgene viral prep # 55636-AAVrg and viral prep # 37825-AAVrg) and with 250 nL of pAAV-hSyn-DIO-hM4D(Gi)-mCherry (Addgene viral prep # 44362-AAV2) or pAAV-hSyn-DIO-mCherry (Addgene viral prep #50459-AAV2) in the LEnt. Following viral vector injections, incisions were sutured, and mice were allowed to recover for 3-4 weeks before the beginning of the behavioral protocol.

### Odor-Taste Sensory Preconditioning Task

Mice were water-deprived the day before starting the behavioral protocol and in the same room where the entire protocol was conducted (Fig. 1A) as previously described^1,8^.

#### Habituation Phase

During habituation, all subjects received 1-hour access to water for three consecutive days in an individual cage to acclimate to the experimental setup.

#### Preconditioning phase

The preconditioning phase consisted of six days of incidental associations (i.e. odor-taste pairings). Each odor-taste pairing was presented on alternating days: on the first day, mice received 1-hour access to a flavored solution consisting of a taste (37.5 mM NaCl; Taste 1, T1) combined with an odorant (0.01% almond; Odor 1, O1) in water, establishing an incidental association between T1 and O1. On the following day, mice were presented with an alternate odor-taste pairing: 0.01% saccharin (Taste 2, T2) combined with 0.05% banana (Odor 2, O2). We repeated this three times as animals received three presentations of each pairing.

#### Conditioning or devaluation phase

This phase involved six days of aversive odor conditioning to create a conditioned stimulus (CS+). On days 1, 3, and 5, mice were given 1-hour access to O1, followed by an i.p. injection of lithium chloride (LiCl, 0.3 M in saline, 1% body weight), which induces gastric malaise, thereby establishing O1 as the conditioned stimulus (CS+). On alternate days (2, 4, and 6), mice received 1-hour access to O2 without the subsequent injection of LiCl, designating O2 as the non-conditioned stimulus (CS-). After this conditioning phase, mice were given a recovery day, during which they had access to water only for 1 hour.

#### Test phase

Over the following two days, mediated and direct learning were assessed in a 1-hour two-choice test. Mediated aversion was evaluated on the first test day, in which mice were given a two-choice test between T1, previously associated with the CS+ but never directly paired with LiCl (referred to as mediated CS+, mCS+), and T2, previously associated with the CS- (mediated CS-, mCS-). On the second test day, direct aversion was assessed by offering a two-choice between CS+ (O1) and CS- (O2).

Data are presented as the percentage of liquid intake for each solution. The total amount of liquid consumption during the two-choice tests and the consumption during preconditioning for the different experimental groups can be found in Supplementary Figures.

### Preference Test

An evaluation of possible innate preferences for odors and tastes was conducted prior to the setup of the odor-taste sensory preconditioning task (Supplementary Figure 1B-C). Mice underwent 3 days of water habituation followed by 6 days of pairings between O1-T1 and O2-T2. After that, mice were subjected to a two-choice bottle test for the evaluation of taste preferences, and on day 11, a two-choice bottle test to check possible odor preferences. Based on these results, the behavioral setting was established between stimuli in which no innate preference was observed.

### Immunohistochemistry and Viral Vectors Signal

To investigate activated areas during our sensory preconditioning protocol, TRAP2:Ai14 mice were perfused 90 min after the start of the test to assess c-Fos expression. Mice were anesthetized with ketamine (100 mg/kg) and xylacine (20 mg/kg) followed by transcardiac perfusion with ice-cold 4% paraformaldehyde (PFA) at pH 7.3. Extracted brains were post-fixed in 4% PFA for 24 hr at 4 ºC and subsequently immersed in 30% sucrose solution until being sectioned on a cryostat (Leica CM3050 S) at 30-μm. Free-floating brain sections were collected and stored in an antifreeze solution at -20 ºC. TdTomato signal was detected in these slices without any further manipulation. For c-Fos immunohistochemistry, sections were washed in 0.1 M phosphate buffer (PB) and incubated for 60 minutes in a blocking solution containing 0.2% Triton X-100 and 5% donkey serum. Sections were then incubated overnight at 4°C with a rabbit anti-c-Fos primary antibody (Synaptic System, #226 008, 1:1000). Following washing, sections were incubated with a donkey anti-rabbit AlexaFluor 488-conjugated secondary antibody (Jackson ImmunoResearch, #711-545-152, 1:1000) for 2 hr at room temperature. After further washes, sections were mounted on glass slides using Fluoromount-G™ medium with DAPI for nuclear staining.

To reveal mCherry viral vector expression in TRAP2:Ai14 mice, brain sections were initially washed, then incubated for 1 hour in a 1% NaOH solution (pH 13) to denature constitutive tdTomato expression, as previously described^29^. Afterward, sections were processed similarly to the c-Fos protocol, with incubation overnight at 4°C using a rat anti-mCherry primary antibody (ThermoFisher, #M11217, 1:500), followed by incubation with a donkey anti-rat AlexaFluor 488-conjugated secondary antibody (Jackson ImmunoResearch, #712-545-150, 1:1000) for 2 hours at room temperature.

To check for viral expression, animals were perfused one week after the end of the behavioral tests. Brain sections were washed to remove antifreeze solution, and thereafter mounted directly to visualize the reporters GFP or mCherry. Animals with minimal or no viral vector expression were excluded from the analysis..

### Image Data Analysis

Fluorescent images were acquired using a Nikon ECLIPSE Ni-E motorized microscope equipped with a 10×/0.40NA objective lens. Image acquisition was conducted via Nis ELEMENTS software and tdTomato, mCherry, GFP, and c-Fos expression were examined in tissue sections across various anterior-posterior coronal levels within the regions of interest.

Quantification of tdTomato and c-Fos-positive cells was performed using ImageJ software version 1.54i (National Institutes of Health, Bethesda, Maryland, USA). A custom macro was developed to detect tdTomato+, c-Fos+, and colocalized+ cells. Cells were classified as tdTomato+ and/or c-Fos+ based on size (diameter approximately 10–15 μm) and fluorescence intensity distinctly above background levels at 10× magnification. For each brain region, 3–5 images per mouse were analyzed, and the mean cell count per mouse was used for graphical representation. The bregma coordinates used for the quantifications in each brain region were as follows: orbitofrontal cortex (OFC; AP +2.80 to +2.46 mm), basolateral amygdala (BLA; AP -1.22 to -1.46 mm), dorsal dentate gyrus (dDG; AP -1.82 to -2.06 mm), and perirhinal cortex (PrH; AP -2.54 to -2.70 mm).

### Statistical Methods

Statistical analyses were conducted using GraphPad Prism 8.0 software. Data are presented as mean ± standard error of the mean (SEM) or before-after for the percentage of liquid consumption. Normality was assessed with the Shapiro-Wilk test. For data following a normal distribution, between-group comparisons were conducted using unpaired or paired t-tests, one-way or two-way ANOVA as appropriate When normality was not met, nonparametric tests were employed (see Supplementary Table for details of sample size and statistics). Outliers, defined as values exceeding two standard deviations above or below the mean of the experimental condition, were identified and removed. A p-value of <0.05 was considered statistically significant.

## Acknowledgements

We would like to thank the personnel of the Animal Facility of the Parc de Recerca Biomedica de Barcelona (PRBB) for mouse care. We thank all the members of our lab for useful discussions during the development of the project. This work was supported by la Generalitat de Catalunya (SGR-00022) from the Departament d’Economia i Coneixement de la Generalitat de Catalunya (Spain) and from the European Research Council (ERC) under the European Union’s Horizon 2020 research and innovation programme (Grant agreement No. [948217]).

## Author contributions

A.B-G. and JA.G.P contributed to the conception of the project and JA.G.P performed and analyzed all experiments. V.A., M.C. and L.V.P. help with different experiments of the project. A.B-G. and JA.G.P wrote the manuscript. All authors have corrected and revised the manuscript.

## Conflict of interest

The authors declare no conflict of interest.

## Notes

### Competing Interest Statement

The authors have declared no competing interest.

### Summary of Updates

We corrected a typo in the name of one author.

