## Supplementary information for "Projecting neurons from lateral entorhinal cortex to basolateral amygdala mediate the encoding of incidental odor-taste associations"

**Content:**

**Supplementary Figures 1-3**

**Supplementary Tables 1-2**

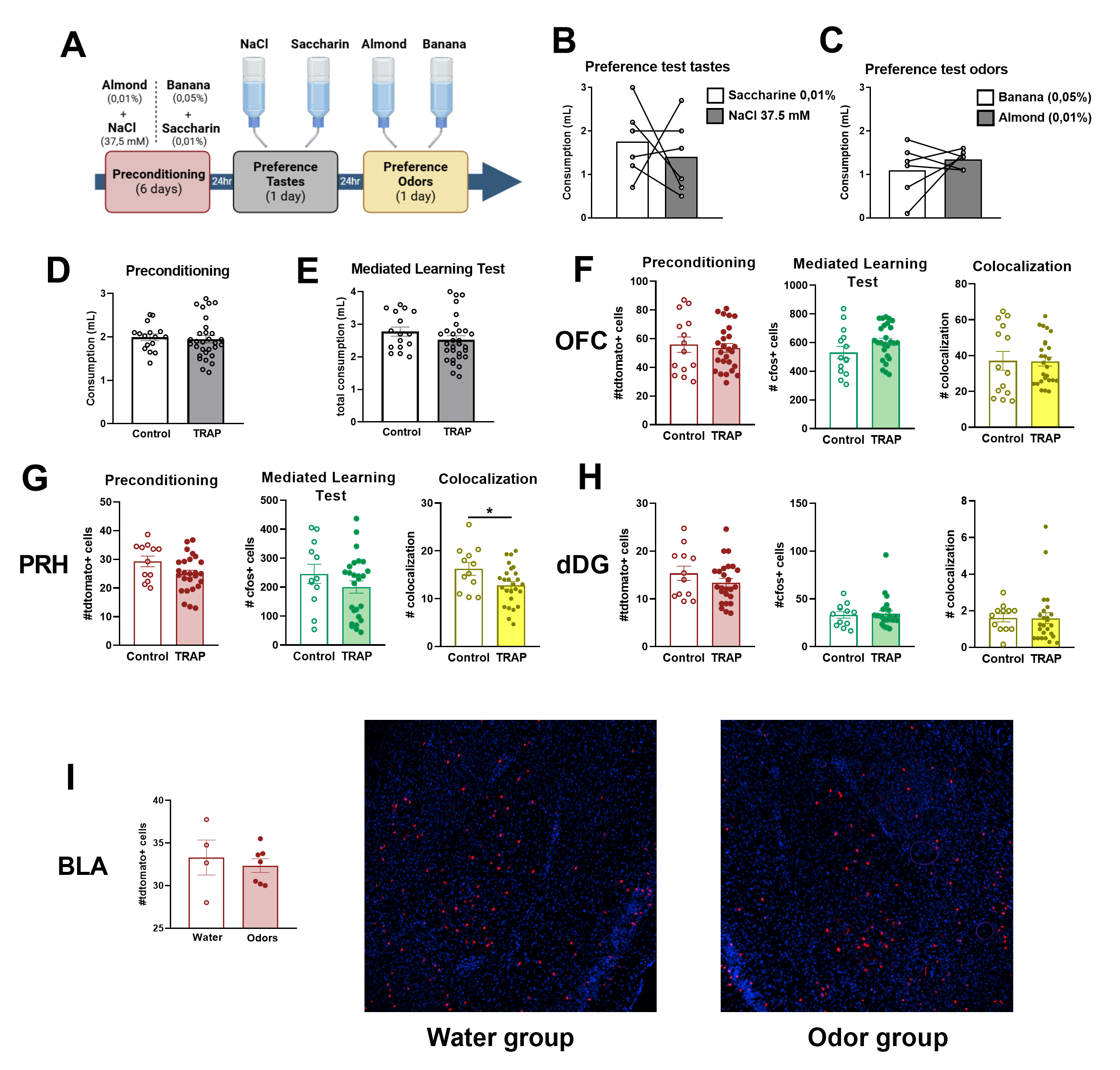
**Fig S1.** **Increase activity of BLA during odor-taste incidental associations**. (A) Schematic illustration of the procedure used to assess potential innate preferences for odors and tastes. Liquid consumption during taste preference test comparing saccharin and NaCl (B) and odor preference test comparing almond and banana (C). Average liquid consumption during the preconditioning phase (D) and total liquid consumption during mediated learning between TRAP and Control groups. Quantification of tdTomato-positive cells, c-Fos-positive cells, and colocalization of tdTomato/c-Fos-positive cells in the orbitofrontal cortex (F), perirhinal cortex (G), and dorsal dentate gyrus (H) of TRAP2:Ai14 mice. (I) Representative images and quantification of tdTomato-positive cells in TRAP2:Ai14 mice exposed to either water or odors. Data are represented as mean ± SEM. For statistical details and n, see Supplementary Table 2. *p<0.05. OFC: Orbitofrontal cortex; PRH: Perirhinal cortex; dDG: dorsal Dentate Gyrus; BLA: Basolateral Amygdala

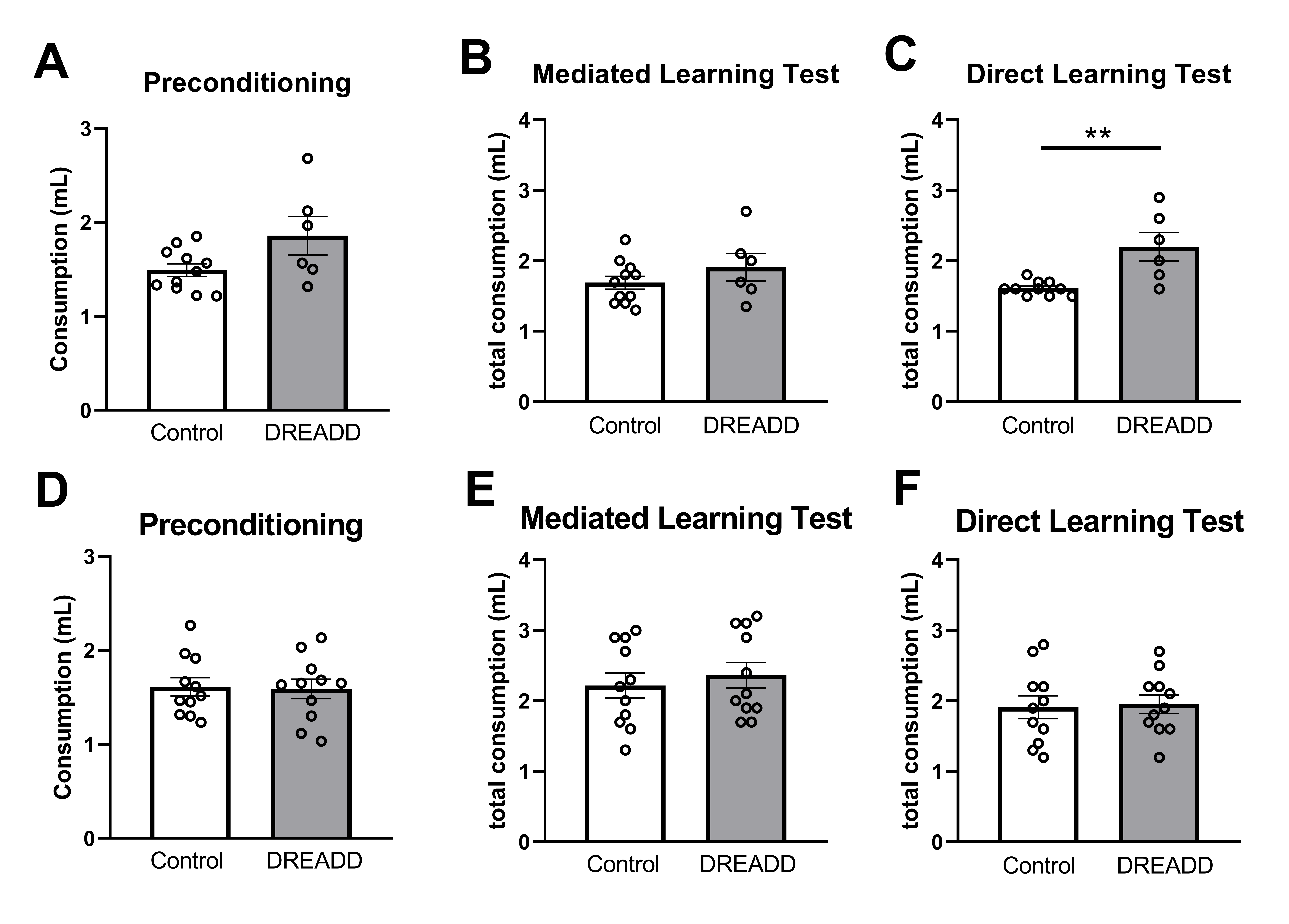

**Fig S2. BLA is necessary for the encoding of incidental associations during odor-taste sensory preconditioning.** (A) Average liquid consumption during the preconditioning phase in Control and DREADD groups injected with J60. (B) Total liquid consumption between groups during mediated learning (B) and direct learning (C) in C57BL/6J mice. Average of liquid consumption between Control and DREADD group during preconditioning (D) and total liquid consumption between groups during mediated learning (E) and direct learning (F) in TRAP2:Ai14 mice. Data are represented as mean ± SEM. For statistical details and n, see Supplementary Table 2. ** p<0.01 (Control vs. DREADD).

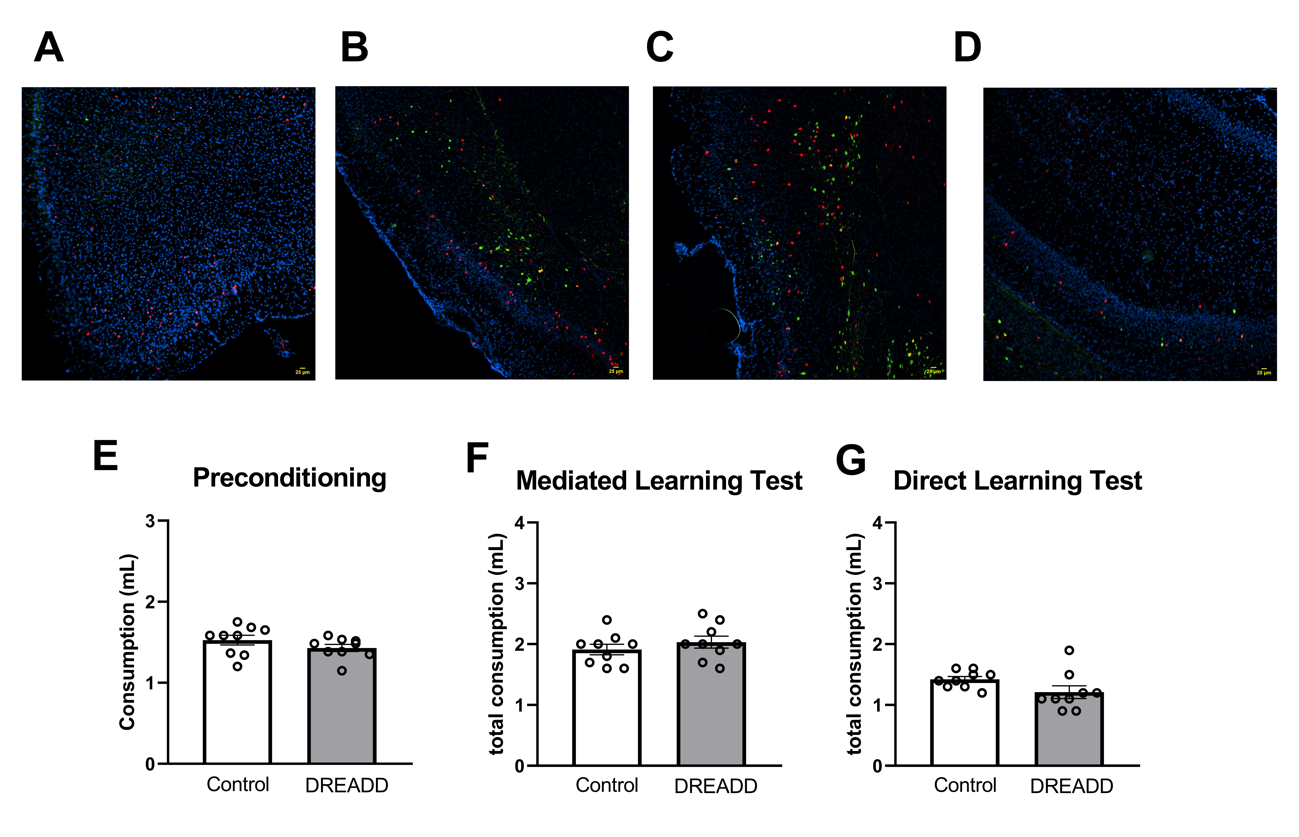

**Fig S3.** **Projections from LEnt to BLA are important for the encoding of incidental associations.** Representative images of activated neurons during incidental associations that project to the basolateral amygdala (BLA) from the orbitofrontal cortex (A), piriform cortex (B), perirhinal cortex (C), and ventral CA1 (D). Average liquid consumption during the preconditioning phase between Control and DREADD groups injected with J60 (E), and total liquid consumption between groups during the mediated learning (F) and direct learning (G) tests in C57BL/6J mice (related to Fig. F-I). Data are presented as mean ± SEM. For statistical details and n, see Supplementary Table 2. Scale bar: 25 μm.

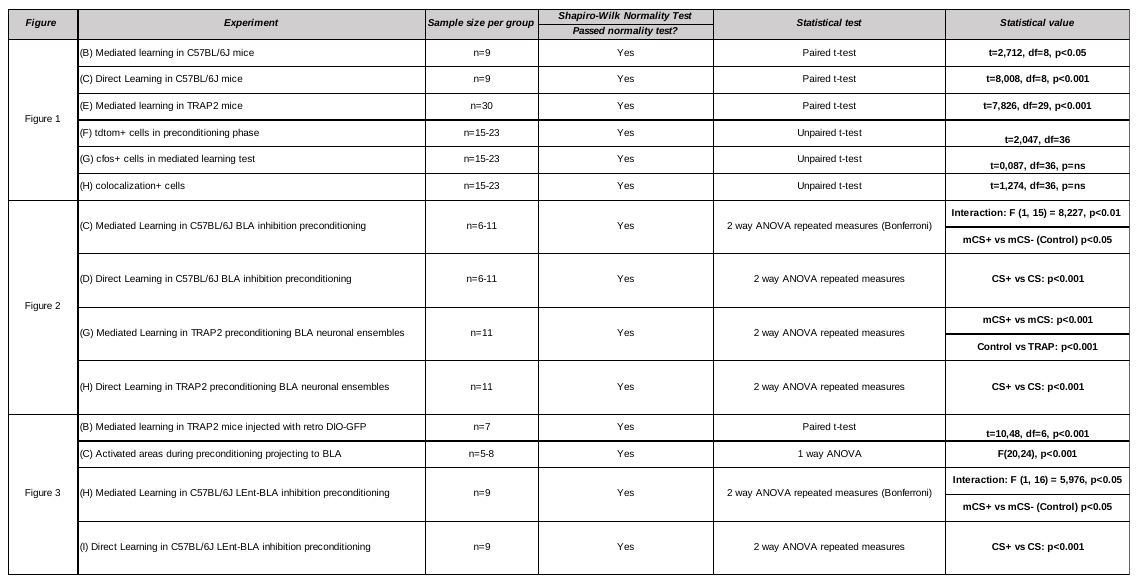
**Table 1.** Statistical analysis. Related to Main Figures 1-3

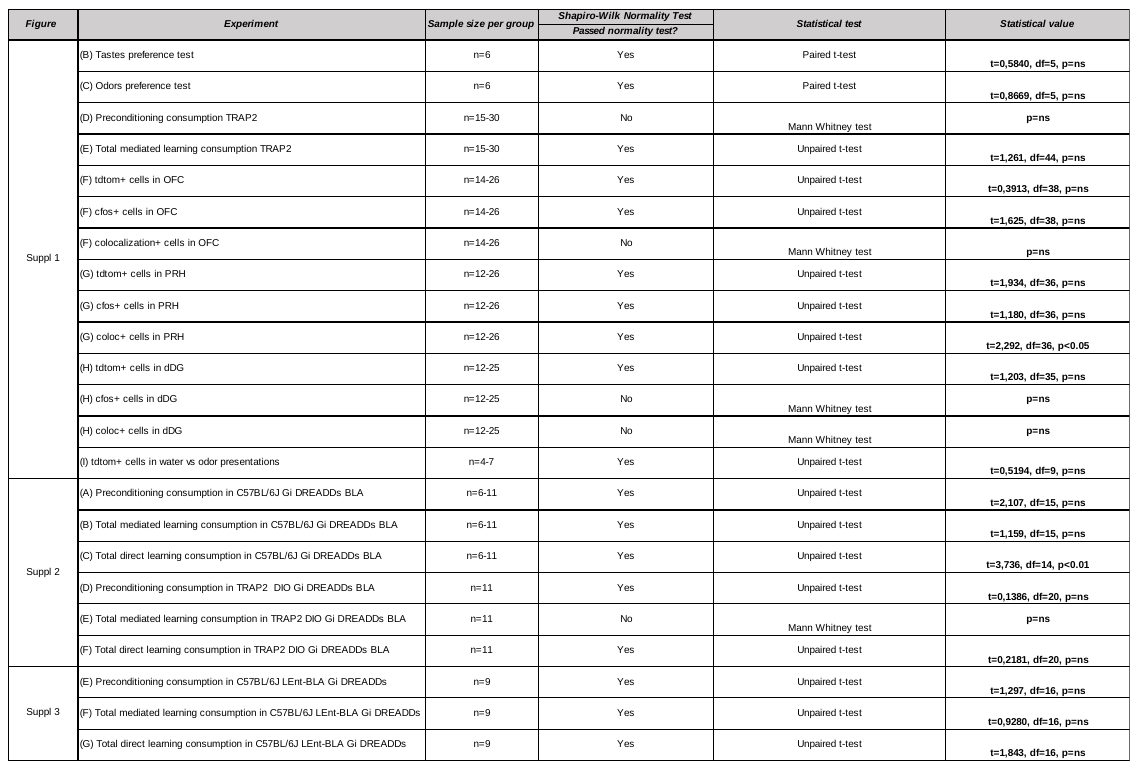

**Table 2.** Statistical analysis. Related to Figures S1-3
